# Bacterial community dynamics during embryonic and larval development of three confamilial echinoids

**DOI:** 10.1101/394478

**Authors:** Tyler J. Carrier, Adam M. Reitzel

## Abstract

Development of some animals is influenced by and, in some cases, dependent on the associated microbiota. The timing of when associated bacterial communities are established during the development of marine invertebrates and their subsequent dynamics across stages are known for only a few species. Here, we compare the bacterial communities of three confamilial echinoids from egg to juvenile using sequence-based approaches. Bacterial communities are established on unfertilized eggs and change gradually during embryonic and larval development. Despite the differences amongst these pre-metamorphic stages, approximately thirty-percent of OTUs identified in association with unfertilized egg were present in the juveniles. During embryonic development, host-associated communities diverged from the environmental microbiota but later converged following the onset of larval feeding. Taken together, the data presented here support the hypothesis that bacterial communities are established prior to fertilization and community composition shifts gradually thereafter, all while remaining distinct from the environment. Future work will need to determine the relative influence of the host and bacteria-bacteria interactions in shaping the associated bacterial community to determine the potential functional importance of bacteria during the development of larval sea urchins and benthic marine invertebrates more broadly.

## INTRODUCTION

The journey from fertilized egg to adulthood has fascinated developmental biologists for centuries. A growing consensus in the field of animal-microbe symbioses suggests that developmental processes are, at least in part, affected by the presence and activity of associated microbiota (McFall-Ngai & Ruby 2000, McFall-Ngai 2002, Fraune & Bosch 2010, Nyholm & McFall-Ngai 2014). The degree of dependency on these communities is highly variable: the removal of bacterial associates inhibits the differentiation of gut tissues in zebrafish (Bates et al. 2006) while other taxa are seemingly unaffected (Manahan et al. 1993). Nevertheless, several patterns have emerged from comparative studies of animal-bacteria interactions. First, animals tend to associate with developmental stage-specific bacterial communities (Apprill et al. 2012, Mortzfeld et al. 2015, Carrier & Reitzel 2018). Second, changes in morphology and life history stage correlate with shifts in the composition and structure of this community (Apprill et al. 2012, Sharp et al. 2012, Mortzfeld et al. 2015, Fieth et al. 2016). Lastly, during ontogeny, animal-associated bacterial communities can be highly specific and diverge from the environmental microbiota through host-mediated selection (Davidson & Stahl 2008, Fraune et al. 2010, Pradea 2011, Franzenburg et al. 2013, McFall-Ngai et al. 2013).

Development of many marine invertebrates includes several morphologically diverse life-history stages. Regardless of taxonomy, reproduction in most species involves an external fertilization, embryogenesis, and larval development before settling and metamorphosing into a benthic juvenile (*e.g*., Thorson 1950, Mileikovsky 1971). When microbial communities are established during early development for marine invertebrates appears to vary among species. Larvae of deep-sea vestimentiferans, for example, are symbiont free until settlement while several species from various taxa are known to vertically inherit microbial symbionts and diversify this community later in development by horizontal (environmental) acquisitions (Nussbaumer et al. 2006, Stewart et al. 2008, Bright & Bulgheresi 2010, Apprill et al. 2012, Sharp et al. 2012, Sipkema et al. 2015). Once established, the location of the bacterial symbionts can also vary. The embryos and larvae of eight echinoderm species have, for example, symbiotic bacteria in the out-pockets of the extracellular matrix surrounding the larval body as well as near or in gut epidermal cells (Cameron & Holland 1983, Bosch 1992, Cerra et al. 1997, Galac et al. 2016, Carrier & Reitzel 2018, Carrier et al. 2018b).

The composition and structure of the bacterial communities associated with echinoderm larvae is highly variable and has been shown to be influenced by food availability (Carrier & Reitzel 2018, Carrier et al. 2018b), phenotype (Carrier & Reitzel 2018), asexual reproduction (Galac et al. 2016), and, potentially, disease (Carrier et al. 2018a). Three confamilial echinoderms (*Strongylocentrotus purpuratus, Mesocentrotus franciscanus*, and *S. droebachiensis*), in particular, associate with species-specific bacterial communities that shift according to larval stage (Carrier & Reitzel 2018). When the diverse and dynamic bacterial community associating with these echinoid larvae are established and how this community varies throughout the entirety of pelagic development is unknown.

By sampling each developmental stage from the unfertilized egg to benthic juvenile and assaying the associated bacterial communities using next-generation sequencing, we test two explicit hypotheses. First, we test if stage-specific bacterial communities during larval development result from associations established in earlier embryonic stages and continue after metamorphosis. Second, we test that three confamilial Strongylocentrotids exhibit relatively similar developmental patterning in the bacterial communities due to their phylogenetic relatedness. Implications from these hypotheses are predicted to complement previous studies showing that the biology of echinoid larvae is inseparable from their microbial community and is an informative system to understand the ecology of animal-microbe symbiosis.

## MATERIALS AND METHODS

### Specimen collection and larval rearing

Adult urchin for each species were collected from populations in the Salish Sea in April 2016 (Carrier & Reitzel 2018). *Strongylocentrotus purpuratus* was hand-collected at low tide at Slip Point, Clallam Bay, WA (48°15′39″ N, 124°15′03″ W; Figure S1) and transferred overnight to the Friday Harbor Laboratories (FHL; University of Washington; Friday Harbor, WA, USA; Figure S1). Similarly, *S. droebachiensis* were hand-collected at low tide at Cattle Point, San Juan Island, WA (48°27′00″ N, 122°57′43″ W; Figure S1) and were transferred to FHL within one hour. *M. franciscanus*, on the other hand, were collected by SCUBA off Bell Island, WA (48°35′49″ N, 122°58′55″ W; Figure S1) and transferred to FHL within two hours. All urchins were suspended in separate sub-tidal cages off the dock at FHL and fed *Nereocystis* spp. (sugar kelp) *ad libitum* until spawning two weeks later.

Urchins were spawned by a one to two mL intracoelomic injection of 0.50 M KCl. Gametes from *S. purpuratus* (n=3 for males and females), *M. franciscanus* (n=1 for males and n=3 for females), and *S. droebachiensis* (n=3 for males and females) were collected, rinsed, and pooled separately for each species. Fertilization of eggs and larval rearing followed Strathmann (1987), with the exception that embryos and larvae were reared using 5.0-μm filtered seawater (FSW) to include the environmental microbiota.

Embryos were incubated in one-liter of FSW at ambient conditions (~10°C and 12°C; Carrier & Reitzel 2018). Two hours post-fertilization, embryos were transferred to triplicate one-liter jars at a density of two individuals•mL^-1^. Beginning at the prism stage, larvae were provided *Rhodomonas lens* at 10,000 cells•mL^-1^, for which was made growth medium-free by centrifugation and resuspension in FSW prior to the introduction to larval cultures. Larval cultures were subsequently diluted to one larvae per two mL at advanced developmental stages. Larval cultures were given 90 to 95% water changes every other day and *R. lens* was replenished at 10,000 cells•mL^-1^.

Monocultures of *R. lens* were grown in f/2 media at room temperature with a combination of ambient and artificial lighting for 24 hours per day (Guillard 1975).

### Sampling across development

Each urchin species was cultured from egg to juvenile. We sampled ten stages: (i) unfertilized eggs, (ii) 16-cell embryo (referred to hereafter as ‘cleavage’), (iii) pre-hatched blastula, (iv) hatched blastula, (v) gastrula, (vi) pre-feeding prism, (vii) 4-, (viii) 6-, and (ix) 8-armed larvae, and (x) post-metamorphic juvenile (see, Table S1 for biological replication for each species at each developmental stage). Developmental stages were identified using a compound microscope (Nikon Eclipse E600; camera: QImaging MicroPublisher 5.0 RTV). At each developmental stage, 100 individuals per jar (n=3) were sampled at random for each species. Immediately after collections, samples were concentrated into a pellet using a microcentrifuge, the FSW was removed with a sterile glass pipette, and pelleted embryos, larvae, or juveniles were then preserved in RNAlater (Thermo Scientific, Massachusetts, USA) and stored at -20°C before DNA extractions.

Complementary to sampling of the developmental stages, the environmental microbiota from the seawater was also sampled. When cultures were sampled, triplicate ~1 L of 5.0-μm FSW was filtered onto a 0.22-μm Millipore filter to retain the environmental microbiota. Full filter disks were then preserved in RNAlater and stored at -20°C before DNA extractions.

### Assaying bacterial communities

Total DNA was extracted from larval samples using the GeneJet Genomic DNA Purification Kit (Thermo Scientific, Massachusetts, USA). For FSW samples, we extracted eDNA using the FastDNA Spin Kit for Soil (MP Biomedical, Illkirch, France). DNA was then quantified using the NanoDrop 2000 UV-Vis Spectrophotometer (Thermo Scientific, Massachusetts, USA) and diluted to 5 ng•μL^-1^ using RNase/DNase-free water.

Bacterial sequences were amplified using universal primers for the V3/V4 regions of the 16S rRNA gene (Forward: 5′ CTACGGGNGGCWGCAG, Reverse: 5′ GACTACHVGGGTATCTAATCC; Klindworth et al. 2013). Products were purified using the Axygen AxyPrep Mag PCR Clean-up Kit (Axygen Scientific, New York, USA), indexed via PCR using the Nextera XT Index Kit V2 (Illumina, California, USA), and then purified again. At each of these three steps, fluorometric quantitation was performed using a Qubit (Life Technologies, California, USA) and libraries were validated using a Bioanalyzer High Sensitivity DNA chip (Agilent Technologies, California, USA). Illumina MiSeq sequencing (v3, 2x300 bp paired-end reads) was performed at the University of North Carolina at Charlotte. PCR recipe and thermal profiles are available in Table S2.

Forward and reverse sequences were paired and trimmed using PEAR (Zhang et al. 2014) and Trimmomatic (Bolger et al. 2014), respectively, converted from fastq to fasta using a custom script, and chimeric sequences were detected using USEARCH (Edgar et al. 2011) and removed using filter_fasta.py prior to the analysis of bacterial 16S rRNA sequences. Using QIIME 1.9.1 (or Quantitative Insights Into Microbial Ecology; Caporaso et al. 2010), bacterial 16S rRNA sequences were analyzed and grouped into operational taxonomic units (OTUs) based on a minimum 97% similarity. The biom table generated by the pick_open_reference_otus.py script was filtered of OTUs with ten or less reads as well as sequences matching the chloroplast of cryptophytes (*i.e., R. lens*; Carrier & Reitzel 2018).

Using the filtered biom table and “biom summarize-table” function to count total sequences per sample, the rarefaction depth of 25,396 was determined and applied to all subsequent analyses (Figure S2). Alpha diversity estimates were calculated using the alpha_diversity.py script and compared statistically by a two-way analysis of variance (ANOVA) in JMP (Pro v. 13). Beta diversity was calculated using unweighted and weighted UniFrac (Lozupone & Knight 2005) as part of jackknifed_beta_diversity.py, and compared using principal coordinate analyses (PCoA) that were then visualized in EMPeror (Vazquez-Baeza et al. 2013) and stylized in Adobe Illustrator CS6. UPGMA-based microbial dendrograms were generated using jackknifed_beta_diversity.py as part of the PCoA analyses. Community similarity across species and developmental stages were compared statistically using an analysis of similarity (ANOSIM) in QIIME as part of the compare_categories.py script, with p<0.05 considered significant. Community composition was generated using summarize_taxa_through_plots.py script and visualized using Prism 7 (GraphPad Software, California, USA). Lastly, observed OTUs were calculated using the alpha_diversity.py script, shared OTUs were calculated using the shared_phylotypes.py script, and the Bray-Curtis index was calculated using the beta_diversity.py script, and compared statistically using ANOVA in JMP (Pro v. 13).

A step-by-step listing of QIIME scripts used to convert raw reads to OTUs for visualization of the data is located in Supplemental Note 1. The raw sequence reads as part of this dataset are available on Dryad.

## RESULTS

### Community-level shifts across development

Urchin-specific bacterial communities (ANOSIM, unweighted: p=0.0011; weighted: p<0.001; Figure 1) shifted significantly in OTU membership (ANOSIM, p=0.0013; Figure 1A; Figures S3–4) and composition (ANOSIM, p=0.0011; Figure 1B; Figures S2–3) during embryonic and larval development. Moreover, community evenness (Shannon and Robbins), diversity (Simpson), richness (Chao1), phylogenetic distance (Faith’s), and Fishers alpha varied significantly across developmental stages (p<0.0001 for all tests) and amongst urchin species (p<0.0001 for all tests) (Figure S5; Table S2).

**Figure 1.**
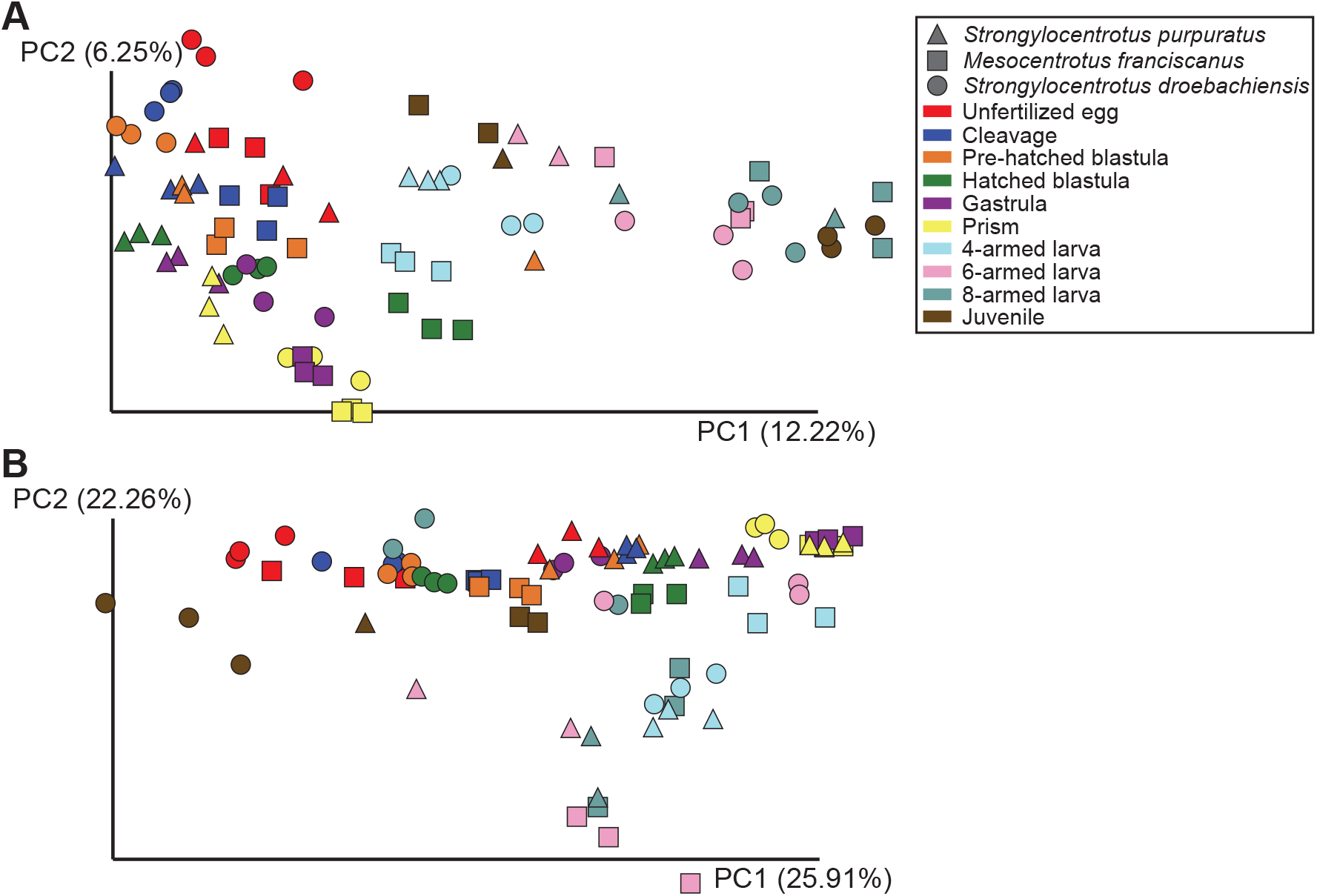
Similarity amongst the associated bacterial community across development for three species of echinoids. Community similarity for *Strongylocentrotus purpuratus* (triangle), *Mesocentrotus franciscanus* (square), and *S. droebachiensis* (circle) for ten developmental stages when considering the presence/absence of taxa (A) and their relative abundance (B).

Unweighted and weighted UniFrac comparisons of the associated bacterial communities for each urchin species suggests that community similarity largely group by developmental stage (Figure 2). Bacterial communities were grouped largely by sequential developmental stages with some specific clusters for: (i) early embryological stages, (ii) late embryogenesis and early larval development, and (iii) later larval stages. The benthic juveniles had more variable positioning on the microbial dendrograms: the bacterial community of *S. purpuratus* juveniles were intermediate to early- and late-stage larvae, *M. franciscanus* juvenile was most similar to eggs, and the *S. droebachiensis* juvenile were most similar to late-stage larvae (Figure 2).

**Figure 2.**
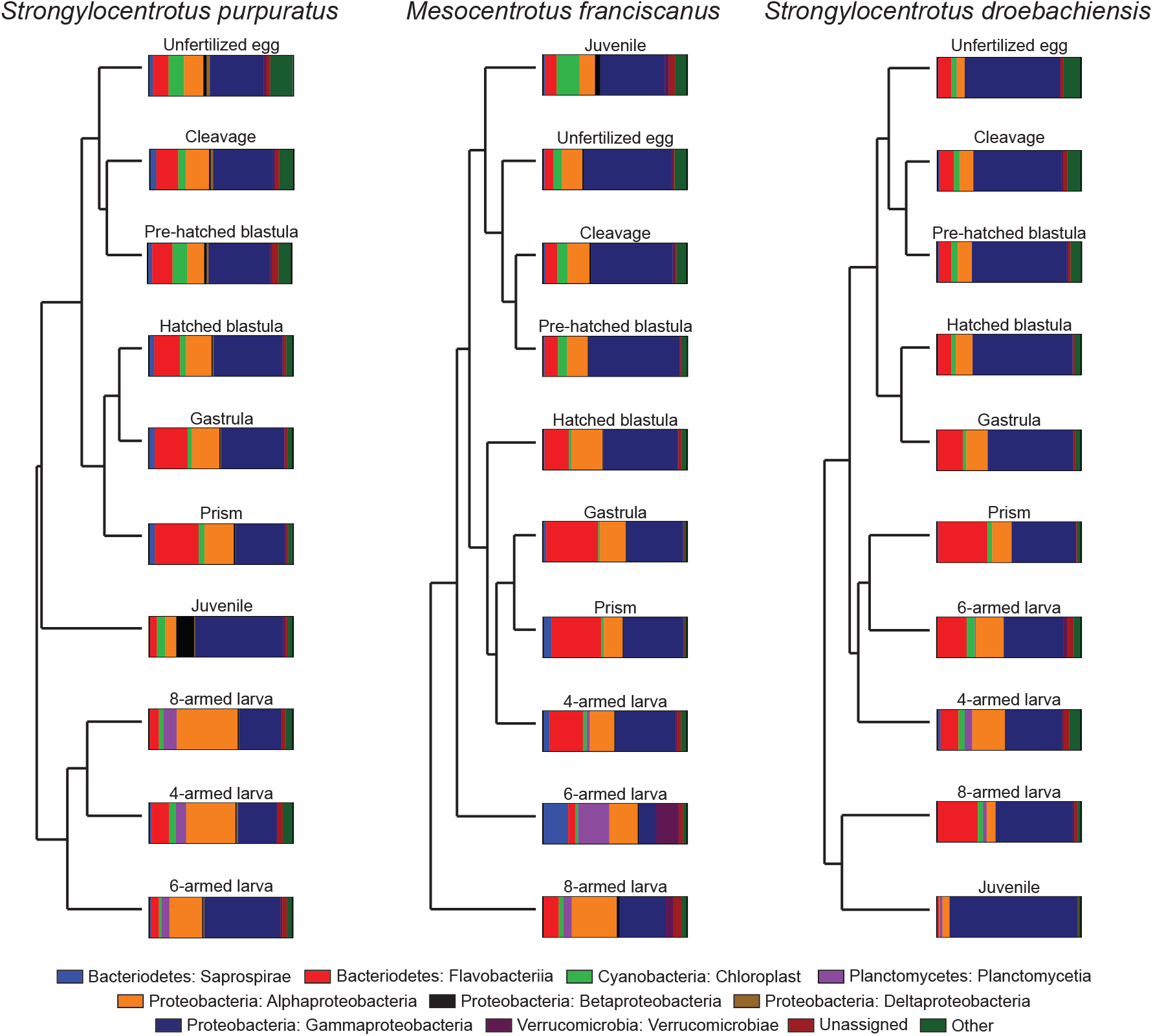
Relatedness of bacterial communities during development. Weighted UniFrac dendrograms and corresponding class-level taxonomic plots for the bacterial communities of *Strongylocentrotus purpuratus* (left), *Mesocentrotus franciscanus* (center), and *S. droebachiensis* (right) across ten developmental stages.

A total of nine classes of bacteria represented at least 1% of the community (Figure 2). Of these, the relative abundance of the γ-proteobacteria (Phylum: Proteobacteria), α-proteobacteria (Phylum: Proteobacteria), and Flavobacteriia (Phylum: Bacteroidetes) were, on average, each more than 10% of the community (Figure 2). For the γ-proteobacteria, the relative abundance ranged from 26.6% to 61.0% for *S. purpuratus*, 12.2% to 62.9% for *M. franciscanus*, and 39.6% to 88.7% for *S. droebachiensis* (Figure 2). Next, for the α-proteobacteria, the relative abundance ranged from 7.7% to 42.4% for *S. purpuratus*, 11.0% to 31.1% for *M. franciscanus*, and 5.1% to 22.7% for *S. droebachiensis* (Figure 2). Lastly, for the Flavobacteriia, the relative abundance ranged from 4.9% to 30.5% for *S. purpuratus*, 5.1% to 36.1% for *M. franciscanus*, and 1.5% to 34.2% for *S. droebachiensis* (Figure 2).

### Dynamics of the associated bacterial species

Total OTUs in association with these confamilial echinoids differed significantly between developmental stages (ANOVA, p<0.0001) for each species (ANOVA, *S. purpuratus*: p<0.0001; *M. franciscanus*: p<0.0001; *S. droebachiensis*: p<0.0001; Figure 3A; Table S1). Unfertilized eggs, on average, associated with a similar number of OTUs amongst species (~1,400 OTUs; ANOVA, p=0.508), which increased during embryonic development (Figure 3A). Following the onset of feeding and larval development, the number of OTUs subsequently decreased until metamorphosis (Figure 3A). The maximum number of associated OTUs during development varied amongst species (*S. purpuratus*: 1,839; *M. franciscanus*: 1,723; *S. droebachiensis*: 1,864) and at which stages this occurred (*S. purpuratus*: gastrula; *M. franciscanus*: 4-armed larva; *S. droebachiensis*: pre-hatched blastula (Figure 3A).

**Figure 3.**
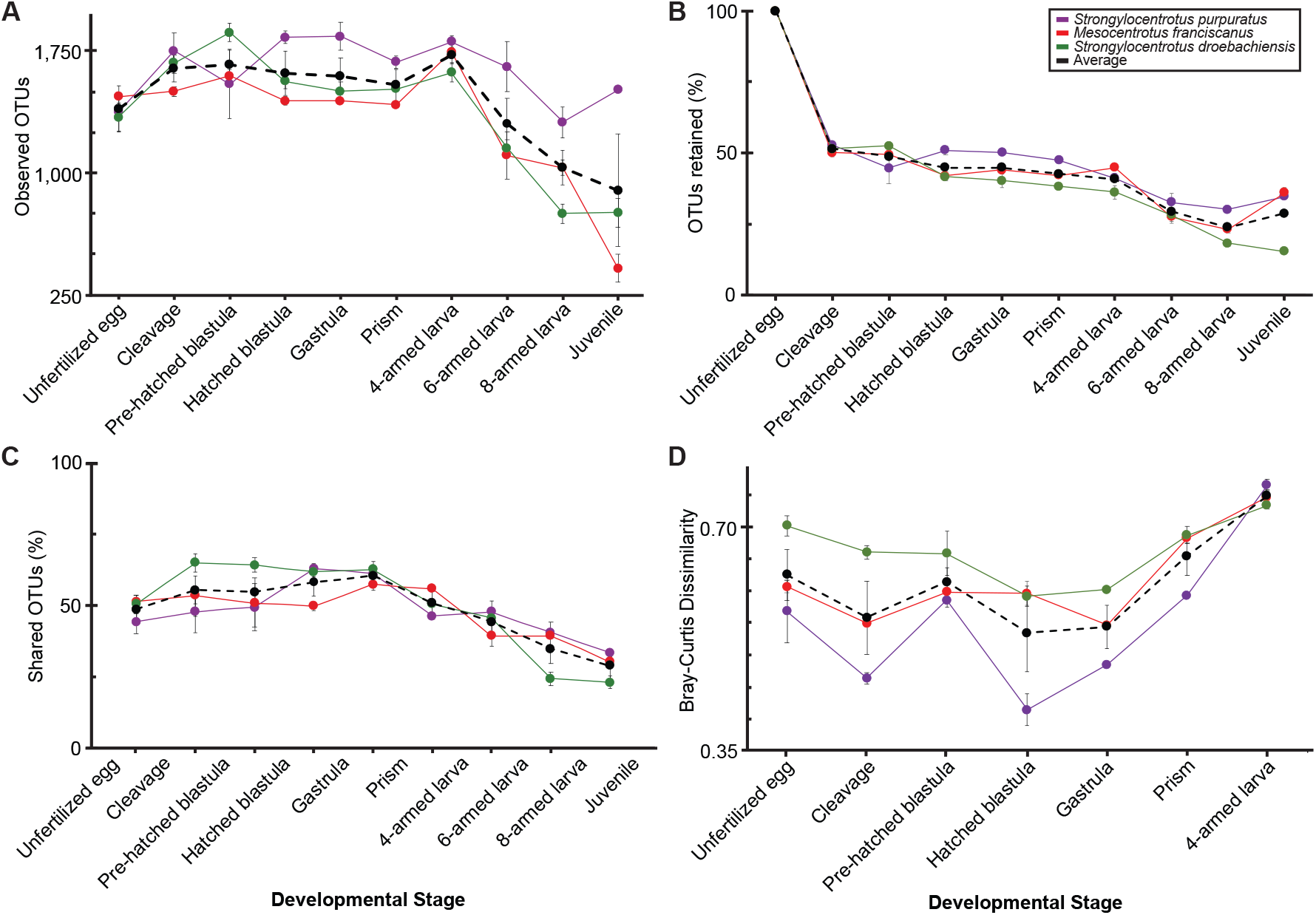
Measures of OTU shifts during development. OTU dynamics across embryonic and larval development for *Strongylocentrotus purpuratus* (purple), *Mesocentrotus franciscanus* (red), and *S. droebachiensis* (green), and the average of the urchin species (black), when considering (A) total observed OTUs, (B) percent of OTUs shared with unfertilized eggs, (C) proportion of OTUs shared between developmental stages, and (D) similarity with the environmental microbiota.

A significant proportion of the OTUs associated with echinoid eggs did not remain in association throughout embryonic and larval development for each urchin species (ANOVA, *S. purpuratus*: p<0.0001; *M. franciscanus*: p<0.0001; *S. droebachiensis*: p<0.0001; Figure 3B; Table S1). Following fertilization and early embryogenesis, ~52% of OTUs from the egg remained in association while ~48% were lost (Figure 3B). In the subsequent larval and juvenile stages, egg-associated OTUs were lost gradually, resulting in juvenile urchins with ~29% of the egg-associated OTUs (Figure 3B). Moreover, when transitioning between any two sequential developmental stages, ~48% of OTUs retained while the rest were specific to either stage (Figure 3C). This result was not simply due to a high diversity of OTUs in the egg samples because OTU number increased after the egg stage for all species (Figure 3A).

Community dissimilarity (Bray-Curtis) between the sea urchin host and environmental microbiota showed significant differences during embryonic and larval developmental for each species (ANOVA, *S. purpuratus*: p<0.0001; *M. franciscanus*: p<0.0001; *S. droebachiensis*: p<0.0001; Figure 3D, Figure S6, Table S1). During development from egg to hatched blastula, the associated bacterial community became more dissimilar from the environmental microbiota (Figure 3D), with the magnitude of dissimilarity varying amongst species (Figure 3D; ANOVA, egg through hatched blastula, each p≤0.0001). Furthermore, following development of the mouth and the onset of feeding, late embryos and early stage larvae became more similar to the environmental bacterial community (Figure 3D). The relative dissimilarity converged in the early larval stages (Figure 3D; ANOVA, 4-armed larvae: p=0.406), which was the last sampled time point for which we determined the bacterial community of the FSW. During divergence and convergence events between the larval host and the environmental microbiota, the environmental bacterial community was consistent throughout the experiment and did not cluster with bacterial communities associated with the urchin (Figure S6).

### *Psychromonas* and echinoid eggs

Bacteria in the genus *Psychromonas* showed the highest abundance for any bacterial genus for each urchin species at any one of the sampled stages, in contrast to their relative rarity (0.7%) in the seawater (Figure 4; Tables S6-7). For each urchin species, the abundance and diversity (Figure S7; ANOVA, p<0.0001 for each) of *Psychromonas* was highest in the unfertilized eggs and declined significantly following fertilization (ANOVA, p<0.0001 for each), until being in very low abundance in 4-armed larvae and the subsequent developmental stages (Figure 4). The magnitude of relative abundance varied amongst urchin species (ANOVA, p<0.0001) and was directly proportion to egg diameter (R^2^=0.833). The OTUs within *Psychromonas* totaled 96 for *S. purpuratus*, 100 for *M. franciscanus*, and 116 for *S. droebachiensis*, while the FSW had 81 (Figure 4). Of those, nine of the ten most abundant *Psychromonas* OTUs for each urchin species were shared amongst urchin species.

**Figure 4.**
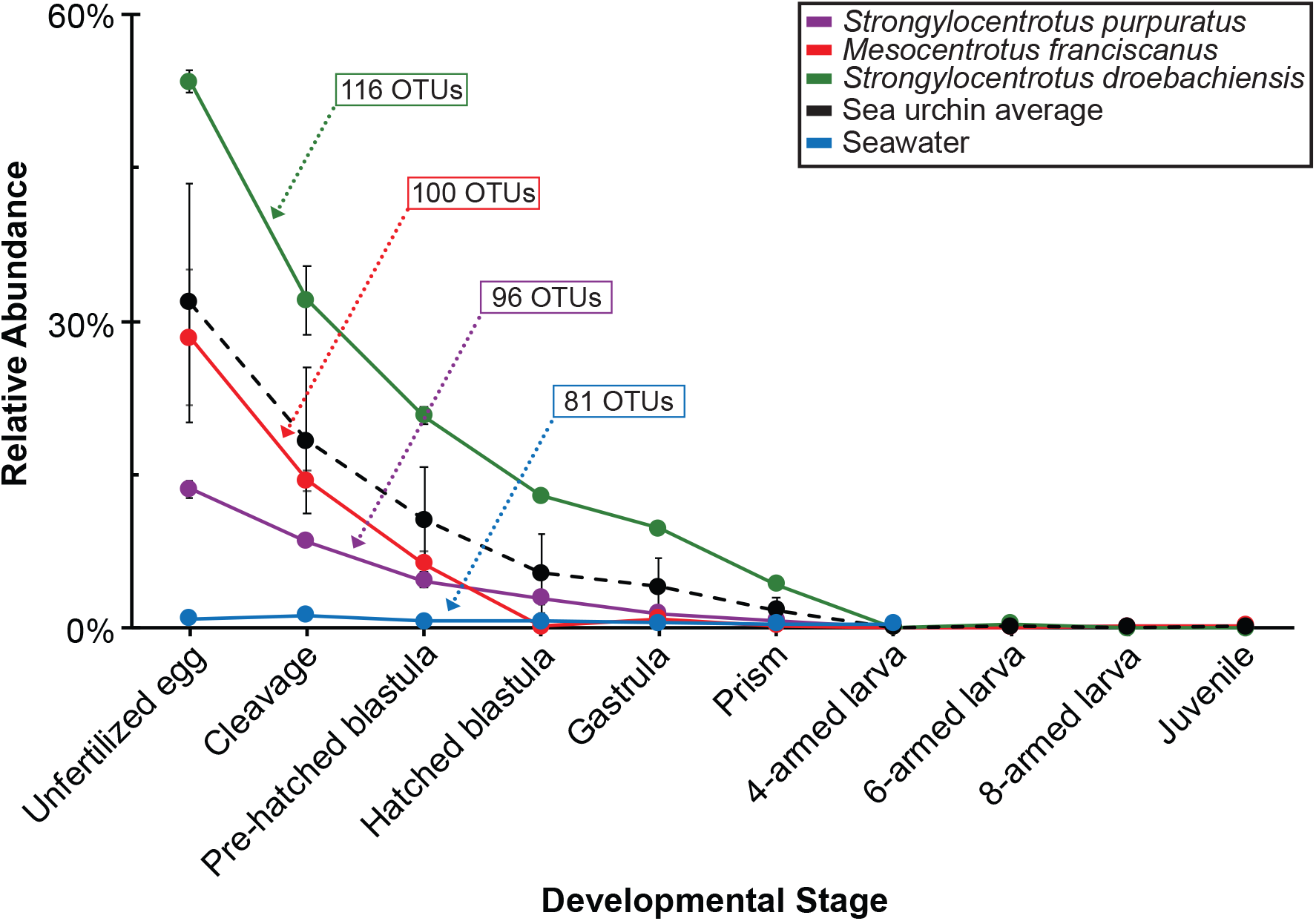
Decay in abundance of *Psychromonas* during development. Relative abundance of the bacterial genus *Psychromonas* across embryonic and larval development for *Strongylocentrotus purpuratus* (purple), *Mesocentrotus franciscanus* (red), *S. droebachiensis* (green), and the seawater (blue) with the number of total OTUs.

## DISCUSSION

Like many marine invertebrates, echinoid life cycles involve a distinct embryonic, larval, and juvenile stage. Our assay of the bacterial communities of three confamilial echinoids supports the hypothesis that the bacterial consortium is highly dynamic across developmental stages, with taxonomic membership and the relative abundance of those taxa shifting gradually from egg to juvenile. Furthermore, over the course of development, the bacterial community of urchin embryos remains distinct and diverges from the environmental microbiota until larvae begin feeding. Lastly, eggs of these echinoids appear to have a specific association with diverse OTUs that match the genus *Psychromonas*.

The bacterial communities associated with echinoplutei are highly dynamic when the larval holobiont faces variation in the abiotic and biotic environment (Carrier & Reitzel 2018, Carrier et al. 2018b). For example, following diet-restriction the bacterial communities of *S. purpuratus, M. franciscanus*, and *S. droebachiensis* each shift considerably from their well-fed counterparts, to potentially cope with reduced exogenous resources (Carrier & Reitzel 2018). When diet-restriction is prolonged and the larval host expresses phenotypic plasticity, the composition and structure of the bacterial community is specific to phenotype, a response correlated with the magnitude in which phenotypic plasticity is expressed (Carrier & Reitzel 2018).

Phenotypic plasticity for echinoid larvae, more broadly, is the interaction between phenotype, nutritional state, and the environment. Characterization of bacterial communities across developmental stages for these three confamilial echinoids extends the understanding of the complex interplay between developmental processes and the surrounding environment in which these species develop. Particularly, after unfertilized eggs establish a bacterial community, the composition of these prokaryotic species shift gradually during embryogenesis and during transitions towards a feeding larva. In these gradual community transitions, it was surprising that more coarse shifts were not observed between major developmental milestones, such as when blastula hatch and shed the fertilization envelope, larvae begin feeding, and/or when larvae metamorphose into benthic juveniles. The latter may be because echinoid larvae undergoing metamorphosis reabsorb some of the larval digestive tract (Chia & Burke 1978). The juveniles therefore may have retained, instead of shed (Fieth et al. 2016), a portion of their bacterial community present in late larval stages.

The influence of particular bacteria and bacterial communities on the development of aquatic and terrestrial invertebrates is well-studied in few species (McFall-Ngai & Ruby 2000, Fraune & Bosch 2010, McFall-Ngai 2014, Nyholm & McFall-Ngai 2014). Established invertebrate systems where bacteria play a significant functional role include *Vibrio fisheri*-mediated maturation of the light organ in the bobtail squid *Euprymna scolopes* (McFall-Ngai & Ruby 2000, McFall-Ngai 2014), immune regulation relating to the epithelial bacteria associated with the freshwater cnidarian *Hydra* spp. (Fraune & Bosch 2010, Bosch 2013, Franzenburg et al. 2013), and the nutritional endosymbionts (e.g., *Buchnera*) of aphids (Douglas 1998). Provided the vast genomic and molecular foundation established over the past several decades for the echinoid clade used in this study (Kudtarkar & Cameron 2017), these confamilial urchins have the potential to be established as a comparative system for beneficial animal-microbe interactions during development (Buckley & Rast 2017, Heyland et al. 2018) and in bi-phasic life-histories. One route for testing the impact of microbial symbionts on echinoid development may be through the egg-associated *Psychromonas*.

Eggs of echinoids have been used as a metric in larval biology, ecology, and evolution (Vance 1973, Wray & Raff 1991, Jaeckle 1995, Moran & McAlister 2009, Marshall et al. 2012) to make inferences about life history mode and transitions (Vance 1973, Collins & Moran 2018), fertilization ecology (Pennington 1985, Levitan 1995), and maternal inheritance (Byrne et al. 1999, Villinski et al. 2002). The high abundance of egg-associated *Psychromonas* and their loss and replacement may imply that this network of bacterial symbionts are functionally connected to early developmental processes.

Interestingly, egg size of echinoids is strongly correlated with total lipid content (Jaeckle 1995, Moran & McAlister 2009) and data here support that egg size is also highly correlated with the abundance of *Psychromonas*. Maternally-inherited lipids for *Dendraster excentricus*, a related echinoid found in the same geographical region of this study, are depleted by the same developmental stage that *Psychromonas* was undetectable here (Rendleman et al. 2018). Moreover, symbiotic bacteria associated with developing embryos of the deep-sea clam *Calyptogena okutanii* is spatiotemporally regulated and has the genetic components to process lipopolysaccharides (Ikuta et al. 2016). *Psychromonas* may, therefore, aid in lipid metabolism (Riley et al. 2008) during embryonic development or serve as a commensal symbiont that relies on the lipids leaching out maturing embryos.

An additional pattern we observed during embryonic development was that urchin-associated bacterial communities diverged from the environmental microbiota. Divergence of host-associated microbiota from the environmental community suggests that urchin embryos and larvae may select, at least, part of their microbial symbiont community. Maintenance of host-specific symbionts may further suggest that these communities are functionally important. To explicitly test how urchin larvae maintain a specific bacterial community and their functional relevance during embryonic and larval development, a multifaceted molecular and -omic (*e.g*., meta-genomic and -transcriptomic) approach would be needed (Williams & Carrier 2018). Approaches for characterizing the mechanisms that establish and regulate host specificity may be adopted from well-established biological systems, such as *Hydra* (Fraune et al. 2010, Fraune & Bosch 2010, Franzenburg et al. 2013) or the winnowing of *Euprymna scolopes* to the luminescent mutualist *Vibrio fisheri* (McFall-Ngai & Ruby 2000, McFall-Ngai 2002, Nyholm & Mcfall-Ngai 2004).

Taken together, data presented here support the hypothesis that the bacterial community associated with the developmental stages of three sea urchin species are established prior to fertilization and shift gradually thereafter. The functional importance and evolutionary significance of this bacterial community and the mechanisms for these corresponding shifts remains unknown. Future experiments involving germ-free culturing as well as manipulations of the host, members of the microbial community, and variable culturing conditions will be needed to determine roles of associated microbiota. Results from these experiments would serve as a key piece towards establishing echinoids as a comparative system for understanding the interdependence of animal development and the associated bacterial communities.

## Acknowledgements

We thank Jason Hodin and Billie Swalla for providing laboratory space; Morgan Eisenlord, Gustav Pauley, Katie Dobkowski, Joe Gaydos, and Richard Strathmann for collecting urchins; Colette Feehan and Richard Strathmann for larval rearing advise; Daniel Janies for sequencing resources; Karen Lopez for technical assistance with sequencing; and the reviewers for improving the specificity and clarity of this manuscript. TJC was supported by an NSF Graduate Research Fellowship, Charles Lambert Memorial Endowment fellowship from the Friday Harbor Laboratories, and a Sigma Xi Grants-in-Aid of Research grant, and AMR was supported by Human Frontier Science Program Award RGY0079/2016.

---

**Figure S1.**
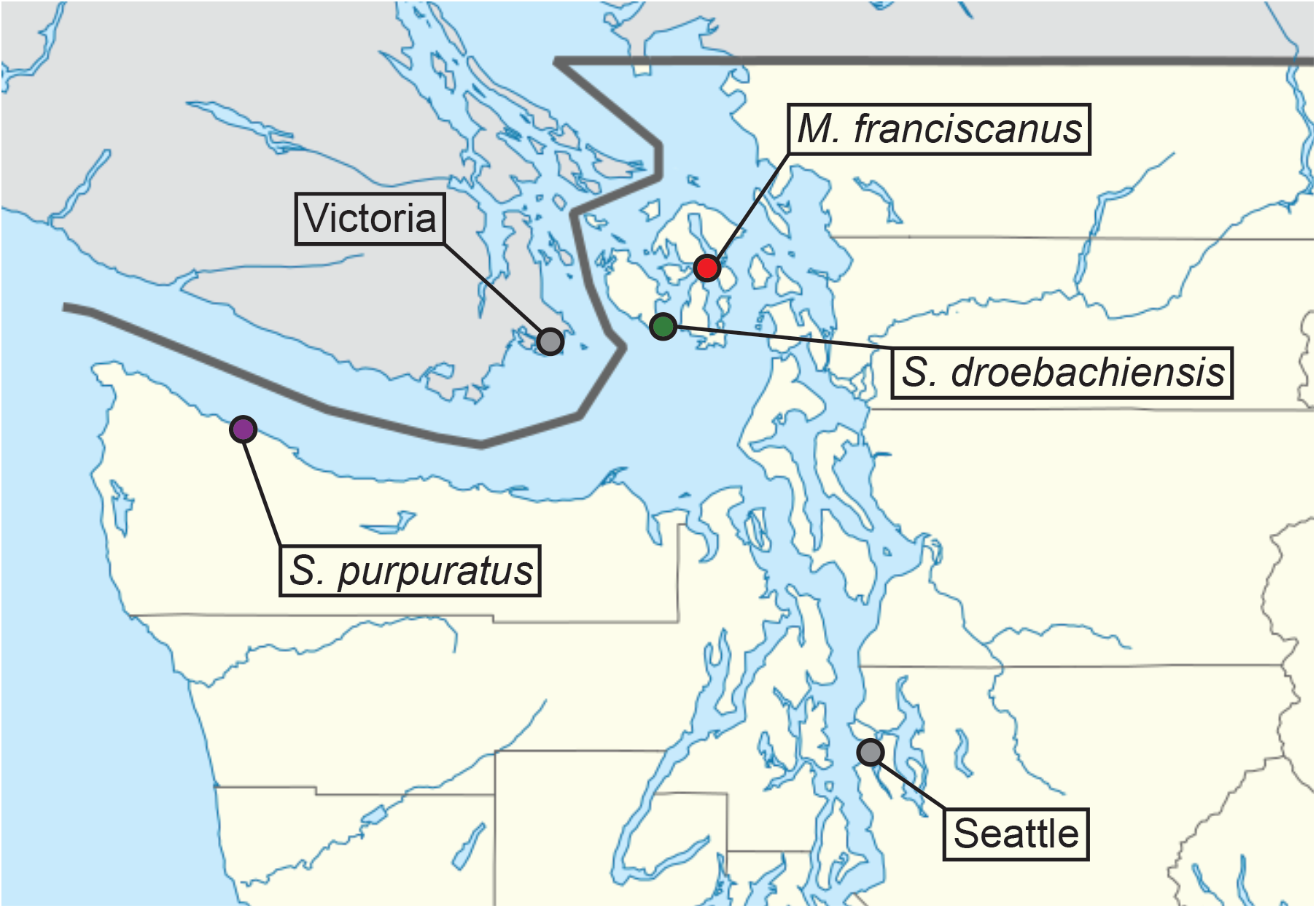
Locations of adult urchin collections throughout the Salish Sea with geographic reference points. Specifically, *Strongylocentrotus purpuratus* were hand-collected at Slip Point, Clallam Bay, WA; *S. droebachiensis* were hand-collected at low tide at Cattle Point, San Juan Island, WA; and, *M. franciscanus* were collected by SCUBA off Bell Island, WA.

**Figure S2.**
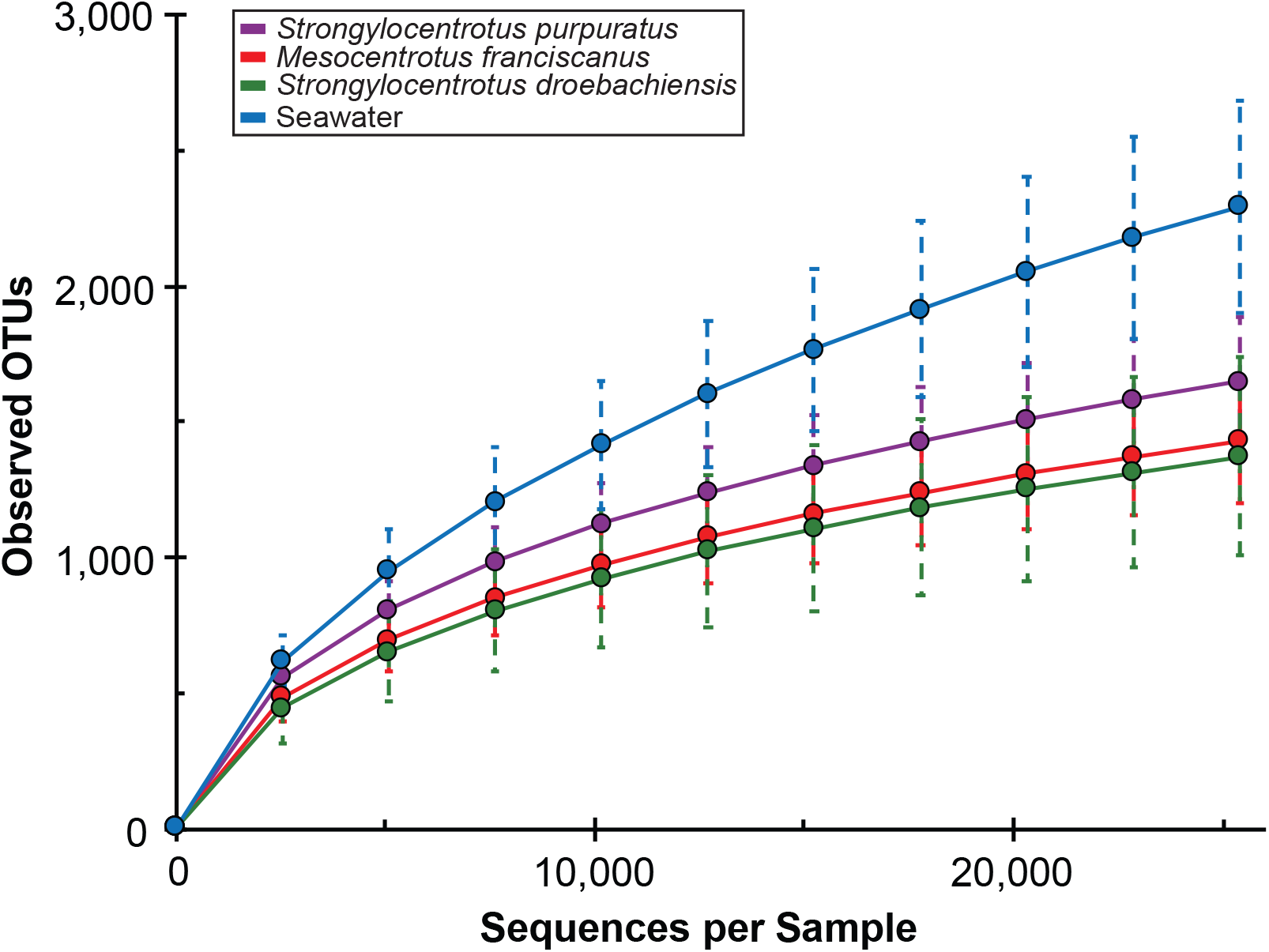
Alpha rarefaction curves for three species of echinoids and of the seawater. Alpha rarefaction curves for *Strongylocentrotus purpuratus* (purple), *Mesocentrotus franciscanus* (red), and *S. droebachiensis* (green) and seawater (blue) based on rarefaction depth of 25,396.

**Figure S3.**
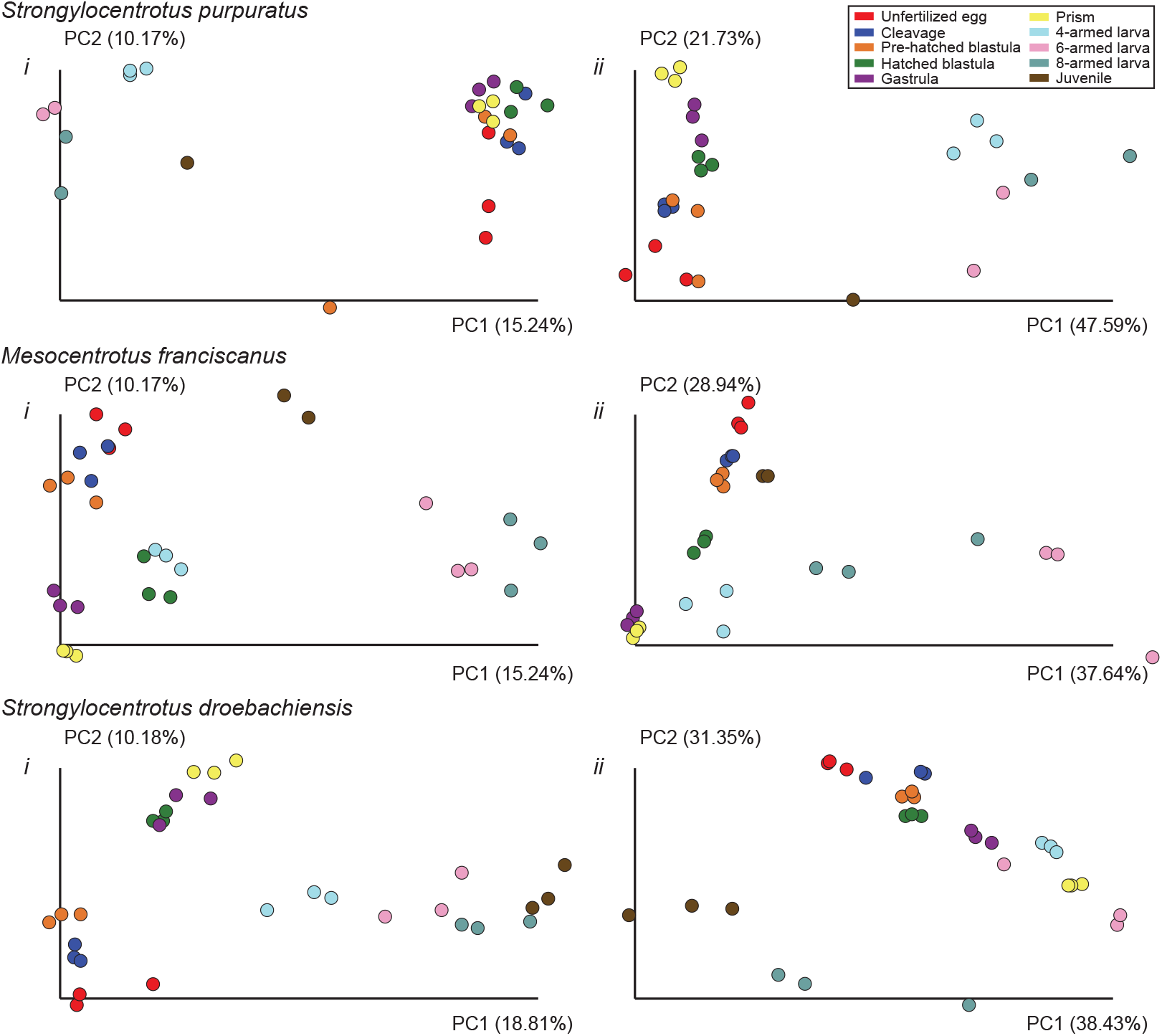
Similarity between the associated bacterial community across development for three species of echinoids. Community similarity for *Strongylocentrotus purpuratus* (top), *Mesocentrotus franciscanus* (middle), and *S. droebachiensis* (bottom) for ten developmental stages when considering the presence/absence of taxa (*i*) and their relative abundance (*ii*).

**Figure S4.**
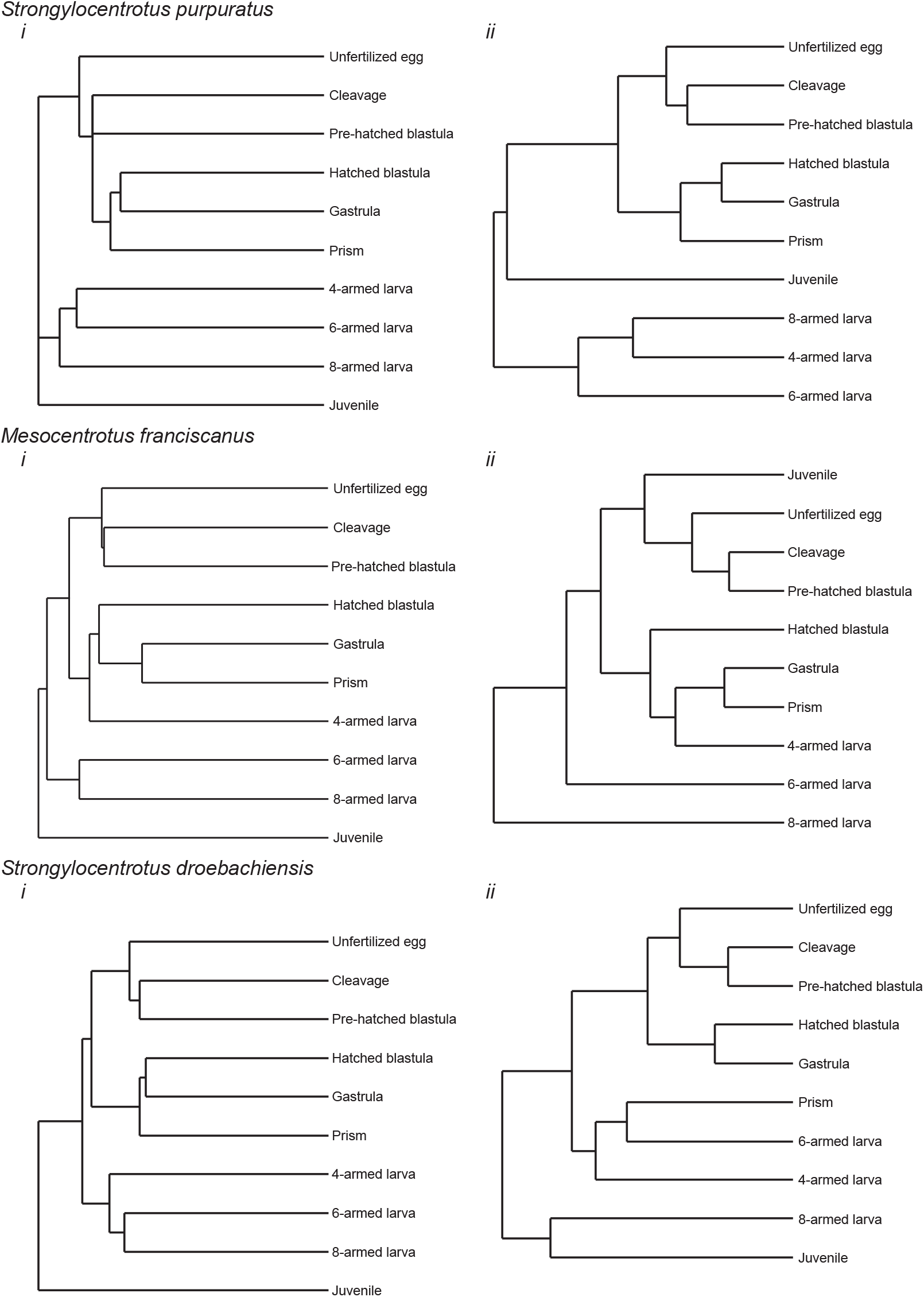
Bacterial community dendrograms across development for three echinoid species. Unweighted (*i*) and weighted (*ii*) bacterial community dendrograms for *Strongylocentrotus purpuratus* (top), *Mesocentrotus franciscanus* (middle), and *S. droebachiensis* (bottom).

**Figure S5.**
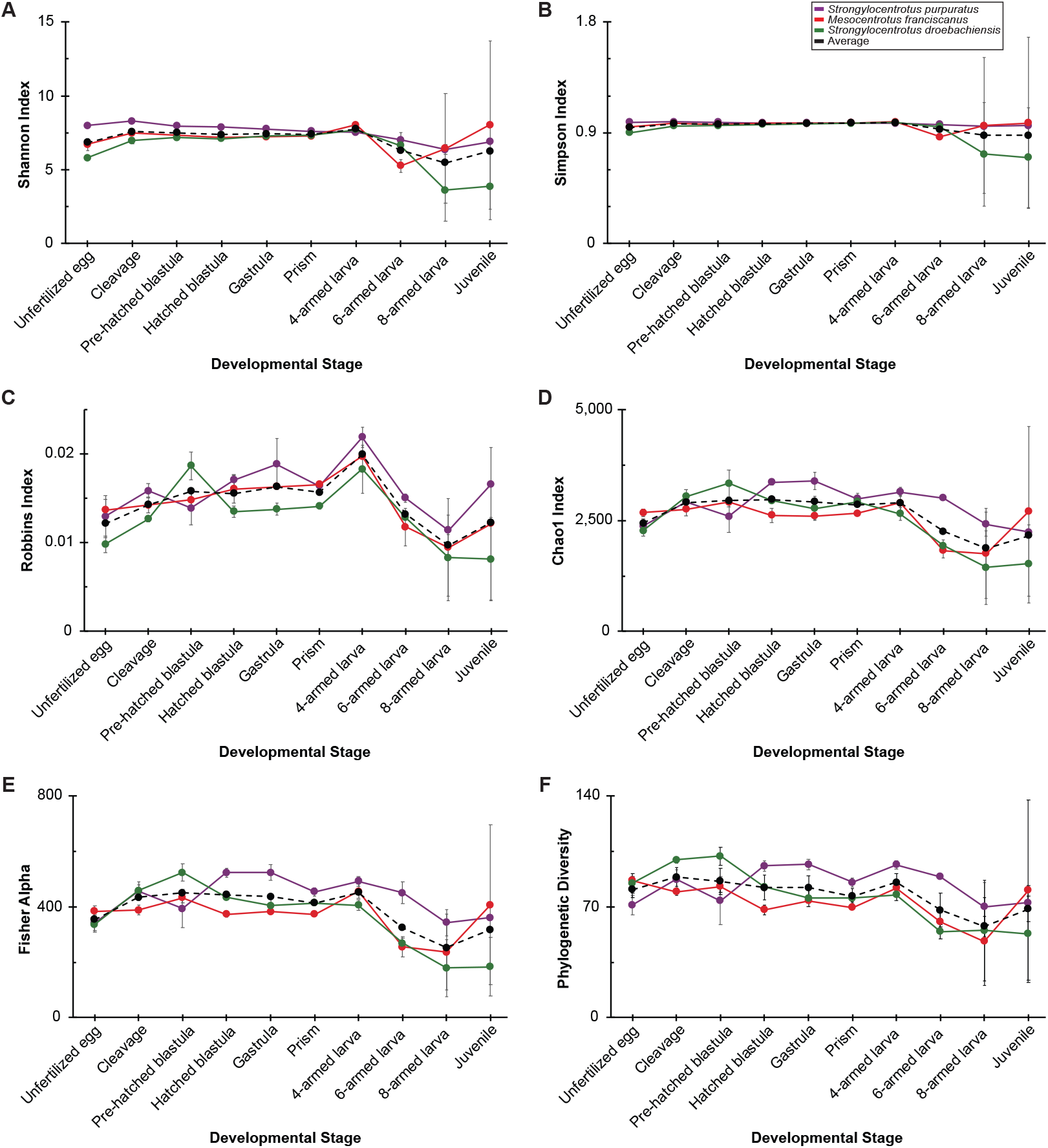
Various alpha diversity indices for three echinoid species across development. Shannon (A), Simpson (B), Robbins (C), Chao1 (D), Fisher (E), and Faiths (F) indices for *Strongylocentrotus purpuratus* (purple), *Mesocentrotus franciscanus* (red), *S. droebachiensis* (green), and the average (black) at ten developmental stages.

**Figure S6.**
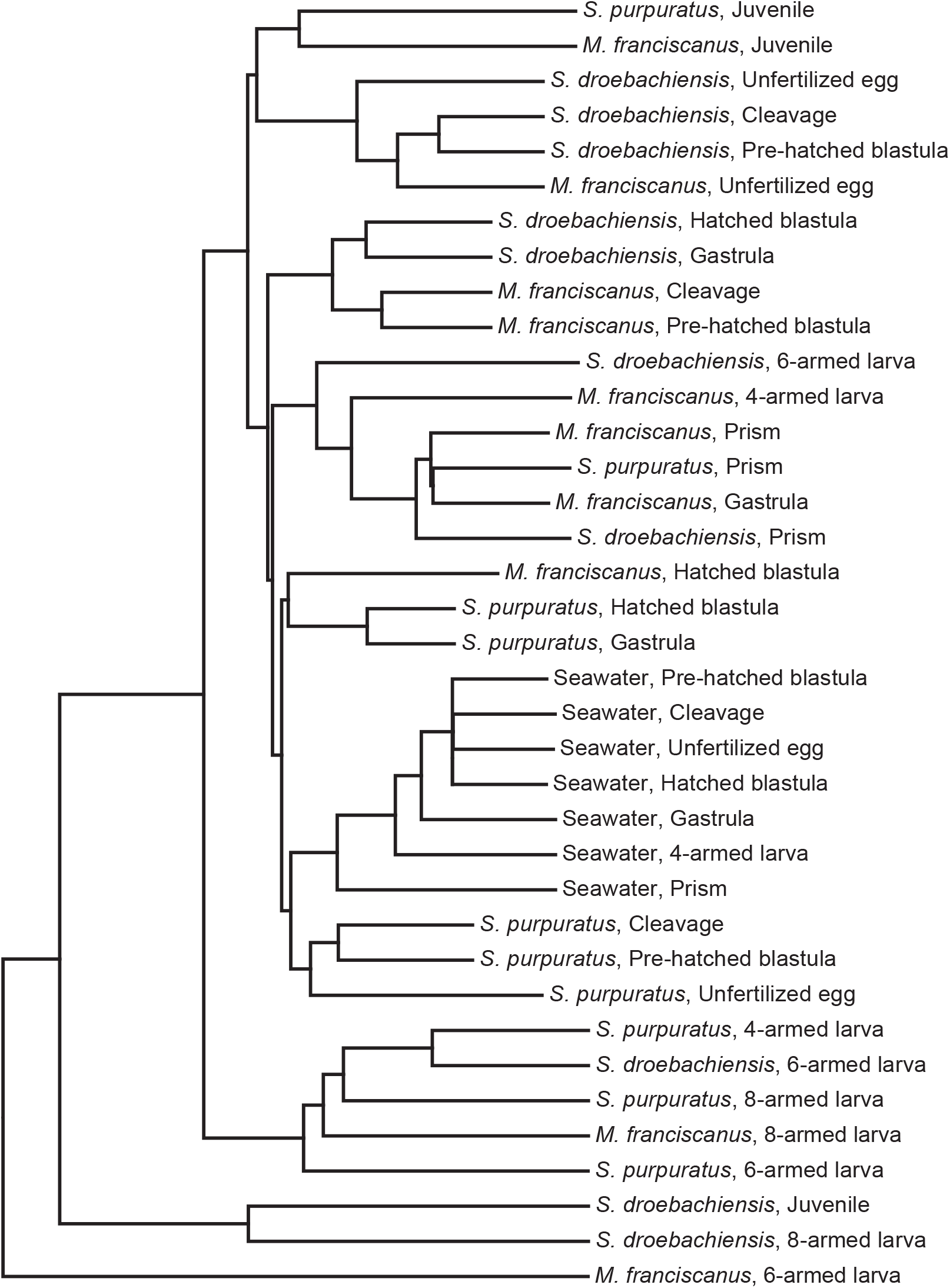
Microbial dendrogram of developmental stages and seawater. Dendrogram of the bacterial communities based on weighted UniFrac for the developmental stages of *Strongylocentrotus purpuratus*, *Mesocentrotus franciscanus*, and *S. droebachiensis* as well as the respective environmental bacterial communities.

**Figure S7.**
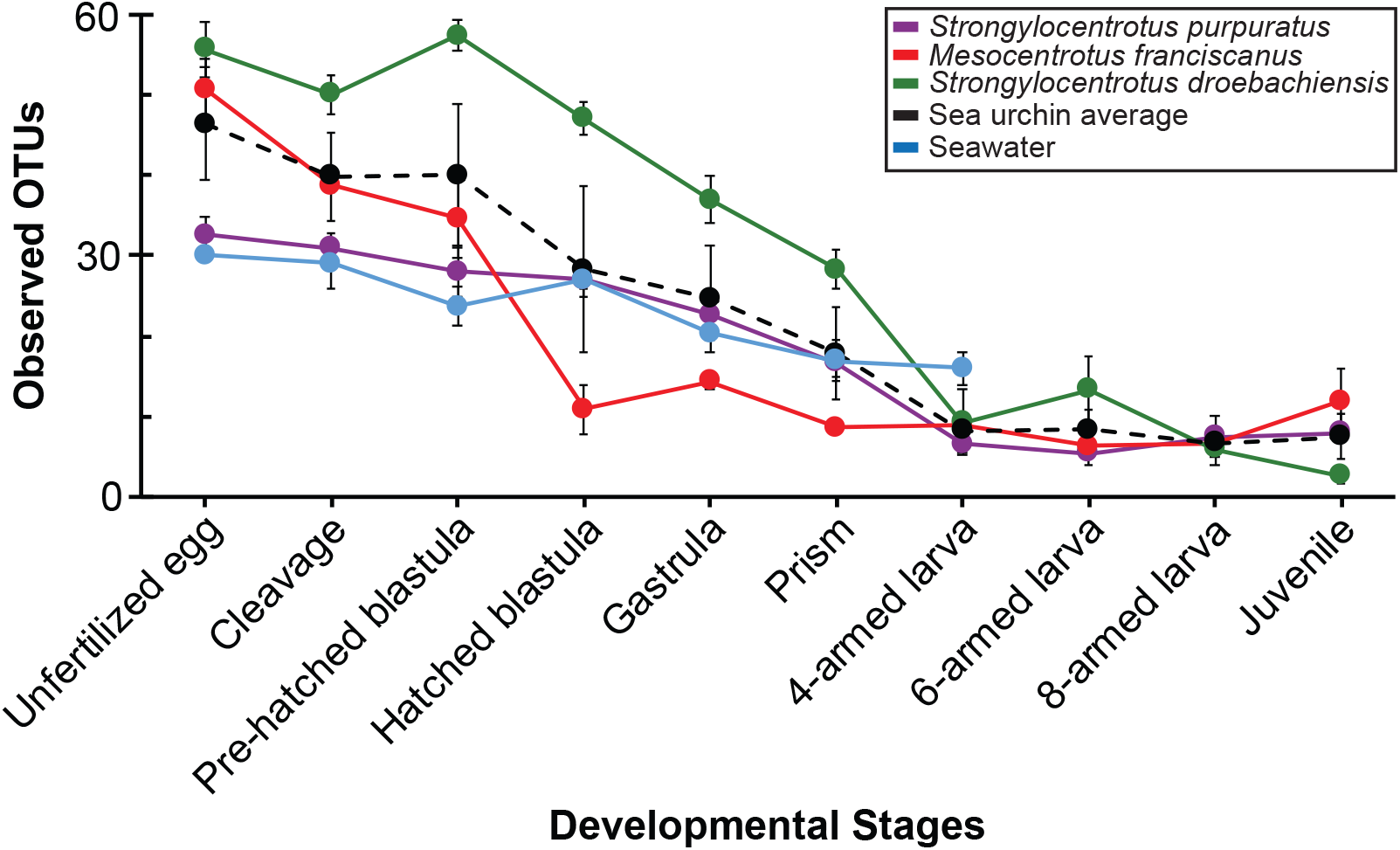
Diversity of *Psychromonas* during development. Total operational taxonomic units (OTUs) within the bacterial genus *Psychromonas* across embryonic and larval development for *Strongylocentrotus purpuratus* (purple), *Mesocentrotus franciscanus* (red), *S. droebachiensis* (green) as well as the average for the three sea urchin species and the in the seawater.

